# A high-throughput neutralizing antibody assay for COVID-19 diagnosis and vaccine evaluation

**DOI:** 10.1101/2020.05.21.109546

**Authors:** Antonio E. Muruato, Camila R. Fontes-Garfias, Ping Ren, Mariano A. Garcia-Blanco, Vineet D. Menachery, Xuping Xie, Pei-Yong Shi

## Abstract

Virus neutralization remains the gold standard for determining antibody efficacy. Therefore, a high-throughput assay to measure SARS-CoV-2 neutralizing antibodies is urgently needed for COVID-19 serodiagnosis, convalescent plasma therapy, and vaccine development. Here we report on a fluorescence-based SARS-CoV-2 neutralization assay that detects SARS-CoV-2 neutralizing antibodies in COVID-19 patient specimens and yields comparable results to plaque reduction neutralizing assay, the gold standard of serological testing. Our approach offers a rapid platform that can be scaled to screen people for antibody protection from COVID-19, a key parameter necessary to safely reopen local communities.

## Text

The ongoing coronavirus disease 2019 (COVID-19) pandemic is caused by severe acute respiratory syndrome coronavirus 2 (SARS-CoV-2), first reported in Wuhan, China in late 2019^1,2^. As of May 18, 2020, COVID-19 has caused 4.8 million confirmed infections and over 318,028 deaths worldwide (https://www.worldometers.info/coronavirus/). Many areas of the world have been in lockdown mode to curb the viral transmission, but the reality is that COVID-19 is here to stay until a safe and efficacious vaccine becomes available. The pandemic’s catastrophic economic impact is pushing governments to reopen their economies, and this creates a public health quandary. At this time, our only option is to minimize viral transmission through social distancing and contact tracing, which relies on the diagnosis of viral RNA through RT-PCR (https://www.fda.gov/media/134922/download). Proper public health policy would be greatly enhanced if we had a reliable and facile assay to measure the immune protection among COVID-19 recovered patients.

Coronavirus infections typically induce neutralizing antibody responses^3^. The seroconversion rates in COVID-19 patients are 50% and 100% on day 7 and 14 post symptom onset, respectively^4^. Given the unknown scale of asymptomatic infections, there is a pressing need for serological diagnosis to determine the real number of infections. Such information is essential for defining the case-fatality rate and for making the policy on the scale and duration of social lockdowns. The serological assays are also required to identify donors with high-titers for convalescent plasma for therapy, and to define correlates of protection from SARS-CoV-2. While viral RNA-based testing for active infection is the current standard, surveying antibody protection is a necessary part of any return to social normality.

For serodiagnosis, several COVID-19 assay platforms have achieved FDA emergency use authorizations (EUA), including ELISA^5^ (https://www.fda.gov/media/137029/download), lateral flow immunoassay (https://www.fda.gov/media/136625/download), and Microsphere Immunoassay (https://www.fda.gov/media/137541/download). These assays measure antibody binding to SARS-CoV-2 spike protein. Since not all spike-binding antibodies can block viral infection, these assay platforms do not functionally measure antibody inhibition of SARS-CoV-2 infection. An ideal serological assay should measure neutralizing antibody levels, which should predict protection from reinfection. Conventionally, neutralizing antibodies are measured by plaque reduction neutralization test (PRNT). Although PRNT and ELISA results generally corelate with each other, the lack of complete fidelity of ELISA continues to make PRNT the gold-standard for determining immune protection^6,7^. However, due to its low throughput, PRNT is not practical for large scale serodiagnosis and vaccine evaluation. This is a major gap for COVID-19 surveillance and vaccine development.

To address the above gap, we developed a fluorescence-based assay that rapidly and reliably measures neutralization of a reporter SARS-CoV-2 by antibodies from patient specimens. The assay was built on a stable mNeonGreen SARS-CoV-2 where the mNeonGreen gene was engineered at the OFR7 of the viral genome^8^. Fig. 1a depicts the flowchart of the reporter neutralization assay in a 96-well format. Briefly, patient sera were serially diluted and incubated with the reporter virus. After incubation at 37°C for 1 h, Vero E6 cells (pre-seeded in a 96-well plate) were infected with the virus/serum mixtures at a multiplicity of infection (MOI) of 0.5. At 16 h post-infection, the mNeonGreen-positive cells were quantitated using a high-content imaging reader (Fig. 1a). Forty COVID-19 serum specimens from RT-PCR-confirmed patients and ten non-COVID-19 serum samples (archived before COVID-19 emergence) were analyzed using the reporter virus. After reporter viral infection, the cells turned green in the absence of serum (Fig. 1b, bottom panel); in contrast, incubation of the reporter virus with COVID-19 patient serum decreased the number of fluorescent cells (top panel). A dose response curve was obtained between the number of fluorescent cells and the fold of serum dilution (Fig. 1c), which allowed for determination of the dilution fold that neutralized 50% of fluorescent cells (NT_50_). The reporter assay rapidly diagnosed fifty specimens in less than 20 h: all forty COVID-19 sera (specimens 1-40) showed positive NT_50_ of 80 to 5152, and all ten non-COVID-19 sera (specimens 41-50) showed negative NT_50_ of <20 for (Fig. 1d).

**Figure 1.**
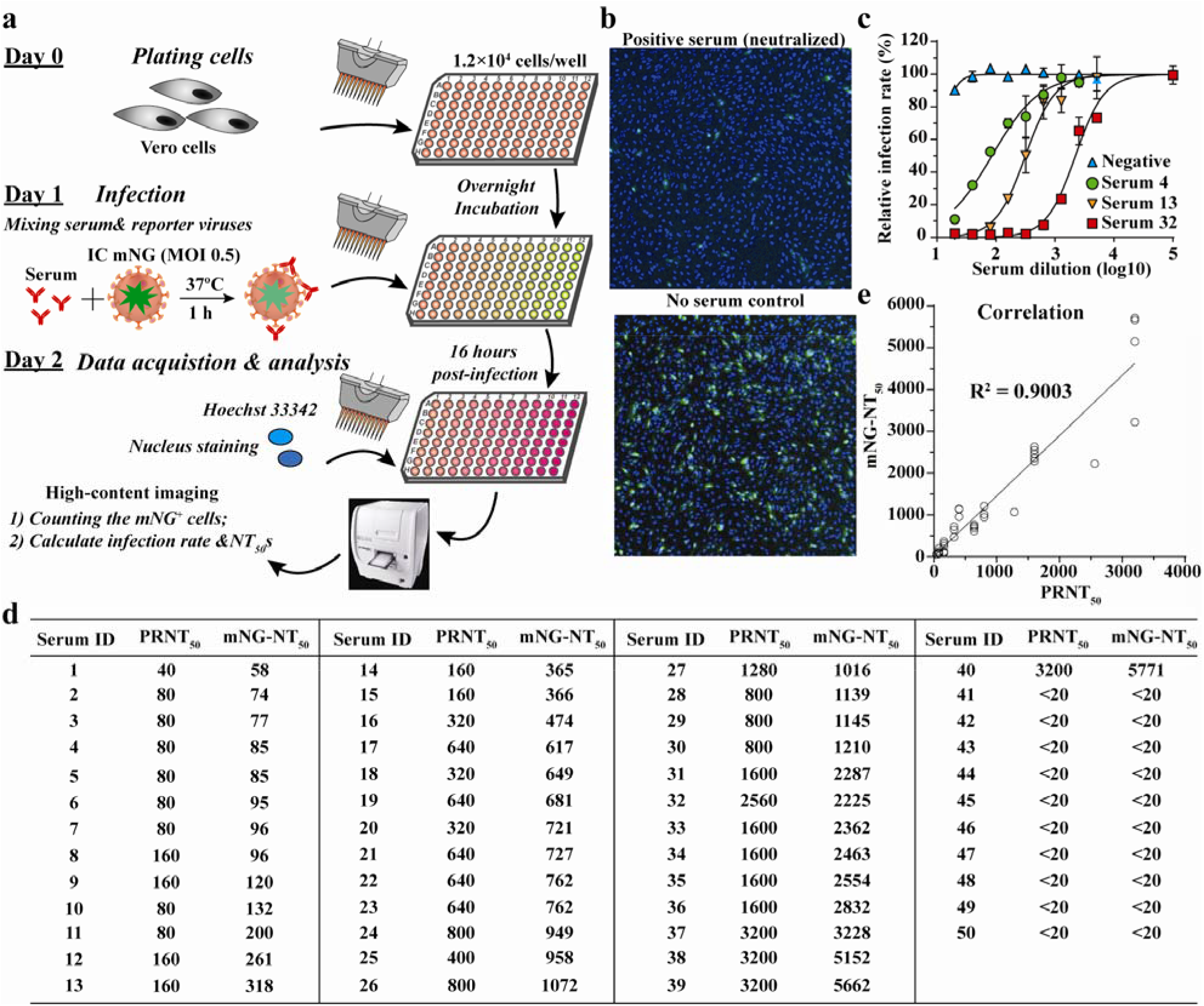
A high-throughput neutralizing antibody assay for COVID-19 diagnosis. (a) Assay flowchart. mNeonGreen SARS-CoV-2 was neutralized with COVID-19 patient sera. Vero E6 cells were infected with the reporter virus/serum mixture with an MOI of 0.5. The fluorescence of infected cells was quantified to estimate the NT_50_ value for each serum. (b) Representative images of reporter virus-infected Vero E6 cells. Images for a positive neutralizing serum (top panel) and no serum control (bottom panel) are presented. (c) Neutralization curves. Representative neutralization curves are presented for three positive sera and one negative sera. (d) Summary of NT_50_ values of fifty patient sera. The NT_50_ values from both reporter virus and conventional PRNT assays are presented. (e) Correlation analysis of NT_50_ values between the reporter virus and PRNT assays. The correlation efficiency *R^2^* is indicated.

To validate the reporter virus neutralization results, we performed the conventional PRNT on the same set of patient specimens. In agreement with the reporter virus results, the forty positive sera showed PRNT_50_ of 40 to 3200, and the ten negative sera exhibited PRNT_50_ of <20 (Fig. 1d). A strong correlation was observed between the reporter virus and PRNT results, with a correlation efficiency *R^2^* of 0.9 (Fig. 1e). The results demonstrate that when diagnosing patient specimens, the reporter virus assay delivers neutralization results comparable to the PRNT assay, the gold standard of serological testing.

Next, we evaluated the specificity of reporter neutralization assay using potentially crossreactive sera and interfering substances (Table 1). Two groups of specimens were tested for cross reactivity. Group I included 138 clinical sera from patients with antigens or antibodies against different viruses, bacteria, and parasites. Group II consisted of 19 samples with albumin, elevated bilirubin, cholesterol, rheumatoid factor, and autoimmune nuclear antibodies. None of the specimens cross neutralized mNeonGreen SARS-CoV-2 (Table 1), including the four common cold coronaviruses (NL63, 229E, OC43, and HUK1). The latter result is consistent with the recent reports that sera from common cold coronavirus patients did not cross react with SARS-CoV-2^5,9^. However, more specimens are required to further validate the cross reactivity, particularly between SARS-CoV-2 and other human coronaviruses, including SARS-CoV-1 and MERS-CoV.

**Table 1.** Cross reactivity of mNeonGreen SARS-CoV-2 neutralization assay

| *Immune sera and #interfering substances | Sample number | Cross reactivity |
| --- | --- | --- |
| Anti-Chikungunya virus | 4 | 0 |
| *Cryptococcus neoformans* antigen | 2 | 0 |
| Anti-Cytomegalovirus | 8 | 0 |
| Anti-Dengue virus | 5 | 0 |
| Anti-Epstein Barr Virus: capsid or nuclear antigen | 8 | 0 |
| Anti-Hepatitis A virus | 5 | 0 |
| Anti-Hepatitis B virus: surface antigen | 14 | 0 |
| Anti-Hepatitis C virus | 3 | 0 |
| Anti-Herpes simplex virus 1 | 7 | 0 |
| Anti-Herpes simplex virus 2 | 5 | 0 |
| Human coronavirus 229E | 1 | 0 |
| Human coronavirus HKU1 | 3 | 0 |
| Human coronavirus NL63 | 1 | 0 |
| Human coronavirus OC43 | 4 | 0 |
| Anti-Human immunodeficiency virus 1 | 7 | 0 |
| Human rhinovirus | 3 | 0 |
| Influenza B virus | 2 | 0 |
| Anti-Measles virus | 7 | 0 |
| Anti-Mumps virus | 5 | 0 |
| Parainfluenza virus 2 | 1 | 0 |
| Parainfluenza virus 4 | 1 | 0 |
| Anti-Parvovirus B19 | 4 | 0 |
| Anti-Rubella virus | 9 | 0 |
| Anti-Syphilis | 4 | 0 |
| Anti-Toxoplasma | 2 | 0 |
| Anti-Typhus Fever | 1 | 0 |
| Varicella zoster virus | 13 | 0 |
| West Nile Virus | 3 | 0 |
| Anti-Yellow fever virus: vaccination | 2 | 0 |
| Anti-Zika virus | 4 | 0 |
| #Albumin (4.5 g/dL) | 3 | 0 |
| #Elevated bilirubin conjugated (>0.4 mg/dL) | 3 | 0 |
| #Elevated bilirubin unconjugated (>0.8 mg/dL) | 3 | 0 |
| #Elevated cholesterol (>200 mg/dL) | 3 | 0 |
| #Elevated rheumatoid factor (>100 IU/mL) | 3 | 0 |
| #Nuclear antibodies | 4 | 0 |

\*A total of 138 sera with antigens or antibodies against different infections (or immunizations) were tested against mNeonGreen SARS-CoV-2 neutralization assay. The immune sera are listed in alphabetical order.
#Tested interfering substances and autoimmune disease nuclear antibodies.

In this study, we developed a rapid fluorescence-based high-throughput assay for COVID-19 serodiagnosis. The reporter virus assay is superior to antigen/antibody binding assays because it measures functional SARS-CoV-2 neutralizing activity in the specimens. When diagnosing patient sera, the reporter virus assay generated NT_50_ values comparable to the conventional PRNT assay. Compared with the PRNT assay, our reporter neutralization test has shortened the assay turnaround time by several days and increased the testing capacity to high throughput. Previously, lentiviruses or vesicular stomatitis virus (VSV) pseudotyped with SARS-CoV-2 spike protein have been reported for neutralization assays^10^. One weakness of the spike pseudotyped assay is that it lacks the same composition of an actual virion, including the SARS-CoV-2 M or E proteins. In addition, the spike protein conformation, either the trimer or monomer, may be different in the pseudotypted virus as compared with the authentic SARS-CoV-2 virion.

Since mNeonGreen SARS-CoV-2 is stable and replicates like wild-type virus, our reporter neutralization assay provides an ideal model for high-throughput serological testing. As the mNeonGreen SARS-CoV-2 grows to >10^7^ PFU/ml in cell culture^8^, the reporter virus can be easily scaled up for testing large sample volumes. Besides mNeonGreen, we have begun to develop other reporter SARS-CoV-2 (*e.g*., luciferase or mCherry) that can also be used for such serological testing. Although the current study performed the assay in a 96-well format, the assay can be readily adapted to 384- and 1536-well formats. Despite the strengths of high throughput and reliability, the current reporter neutralization assay must be performed in biosafety level 3 (BSL3) containment. Efforts are ongoing to engineer an attenuated version of SARS-CoV-2 so that the assay could be performed at a BSL2 facility. Nevertheless, the mNeonGreen reporter assay offers a rapid, high-throughput platform to test COVID-19 patient sera not previously available.

Because neutralizing titer is a key parameter to predict immunity, the reporter neutralization assay should be useful for high-throughput evaluation of COVID-19 vaccines and for identification of high neutralizing convalescent plasma for therapy. Indeed, treatment of severe COVID-19 patients with convalescent plasma shows clinical benefits^11^. For vaccine development, a standardized neutralizing assay will facilitate down selection of various candidates for clinical development. Furthermore, the reporter assay could be used over time to monitor the waning of protective neutralizing titers in COVID-19 patients and to study the correlates of protection from SARS-CoV-2. Thus, the ability to rapidly measure neutralizing antibody levels in populations is essential for guiding policymakers to reopen the economy and society, deploy healthcare workers, and prepare for SARS-CoV-2 reemergence.

## Methods

### mNeonGreen SARS-CoV-2

The virus stock of mNeonGreen SARS-CoV-2 was produced using an infectious cDNA clone of SARS-CoV-2 in which the ORF7 of the viral genome was replaced with reporter mNeonGreen gene^8^. After rescued from the genome-length viral RNA-electroporated cells, the viral stock was prepared by amplifying the mNeonGreen SARS-CoV-2 on Vero E6 cells for one or two rounds. The titer of the virus stock was determined by a standard plaque assay.

### Human sera and interfering substances

All suman serum specimens were obtained at the University of Texas Medical Branch (UTMB). All specimens were de-identified from patient information. A total of forty de-identified convalescent sera from COVID-19 patients (confirmed with viral RT-PCR positive) were tested in this study. Ten non-COVID-19 sera, collected before COVID-19 emergence^12,13^, were also tested in the reporter virus and PRNT assays. For testing cross reactivity, a total of 138 de-identified specimens from patients with antigens or antibodies against different viruses, bacteria, and parasites were tested in the mNeonGreen SARS-COV-2 neutralization assay (Table 1). For testing interfering substances, nineteen de-identified serum specimens with albumin, elevated bilirubin, cholesterol, rheumatoid factor, and autoimmune nuclear antibodies were tested in the reporter neutralization assay. All human sera were heat-inactivated at 56°C for 30 min before testing.

### mNeonGreen SARS-CoV-2 reporter neutralization assay

Vero E6 cells (1.2×10^4^) in 50 μl of DMEM (Gibco) containing 2% FBS (Hyclone) and 100 U/ml Penicillium-Streptomycin (P/S; Gibco) were seeded in each well of black μCLEAR flat-bottom 96-well plate (Greiner Bio-one™). The cells were incubated overnight at 37°C with 5% CO_2_. On the following day, each serum was 2-fold serially diluted in 2% FBS and 100 U/ml P/S DMEM, and incubated with mNeonGreen SARS-CoV-2 at 37°C for 1 h. The virus-serum mixture was transferred to the Vero E6 cell plate with the final multiplicity of infection (MOI) of 0.5. For each serum, the starting dilution was 1/20 with nine 2-fold dilutions to the final dilution of 1/5120. After incubating the infected cells at 37°C for 16 h, 25 μl of Hoechst 33342 Solution (400-fold diluted in Hank’s Balanced Salt Solution; Gibco) were added to each well to stain cell nucleus. The plate was sealed with Breath-Easy sealing membrane (Diversified Biotech), incubated at 37°C for 20 min, and quantified for mNeonGreen fluorescence on Cytation™ 7 (BioTek). The raw images (2×2 montage) were acquired using 4× objective, processed, and stitched using the default setting. The total cells (indicated by nucleus staining) and mNeonGreen-positive cells were quantified for each well. Infection rates were determined by dividing the mNeonGreen-positive cell number to total cell number. Relative infection rates were obtained by normalizing the infection rates of serum-treated groups to those of non-serum-treated controls. The curves of the relative infection rates versus the serum dilutions (log10 values) were plotted using Prism 8 (GraphPad). A nonlinear regression method was used to determine the dilution fold that neutralized 50% of mNeonGreen fluorescence (NT_50_). Each serum was tested in duplicates. All mNeonGreen SARS-CoV-2 reporter neutralization assay was performed at the BSL-3 facility at UTMB.

### Plaque reduction neutralization test (PRNT)

Vero E6 cells (1.2×10^6^ per well) were seeded to 6-well plates. On the following day, 100 PFU of infectious clone-derived wild-type SARS-CoV-2 was incubated with serially diluted serum (total volume of 200 μl) at 37°C for 1 h. The virus-serum mixture was added to the pre-seeded Vero E6 cells. After 1 h 37°C incubation, 2 ml of 2% high gel temperature agar (SeaKem) in DMEM containing 5% FBS and 1% P/S was added to the infected cells. After 2 days of incubation, 2 ml neutral red (1 g/l in PBS; Sigma) was added to the agar-covered cells. After another 5-h incubation, neutral red was removed. Plaques were counted for NT_50_ calculation. Each serum was tested in duplicates. The PRNT assay was performed at the BSL-3 facility at UTMB.

### Statistical analysis

The correlation of the NT_50_ values from mNeonGreen reporter SARS-CoV-2 assay and the PRNT_50_ values from plaque neutralization assay was analyzed using a linear regression model in the software Prism 8 (GraphPad).

### Data availability

The results presented in the study are available upon request from the corresponding authors. The mNeonGreen reporter SARS-CoV-2 has been deposited to the World Reference Center for Emerging Viruses and Arboviruses (https://www.utmb.edu/wrceva) at UTMB for distribution.

## Acknowledgements

We thank colleagues at UTMB for helpful discussion during the course of this project. P.-Y.S. was supported by NIH grants AI142759, AI134907, AI145617, and UL1TR001439, and awards from the Sealy & Smith Foundation, Kleberg Foundation, John S. Dunn Foundation, Amon G. Carter Foundation, Gilson Longenbaugh Foundation, and Summerfield Robert Foundation. M.A.G.-B. was supported by NIH grant CA204806 and the Vacek Distinguished Chair. V.D.M. was supported by NIH grants U19AI100625, R00AG049092, R24AI120942, and STARs Award from the University of Texas System. A.E.M. is supported by a Clinical and Translational Science Award NRSA (TL1) Training Core (TL1TR001440) from NIH. C.R.F.-G. is supported by the predoctoral fellowship from the McLaughlin Fellowship Endowment at UTMB.

## Author contributions

P.R., M.A.G.-B., V.D.M., X.X., and P.-Y.S conceived the study. A.E.M. and C.R.F.-G. performed the experiments and analyzed the results. P.R. prepared the serum specimens. M.A.G.-B., V.D.M., X.X., and P.-Y.S wrote the manuscript.

